# Single-Cell Analysis Reveals Tissue-Specific T Cell Adaptation and Clonal Distribution Across the Human Gut-Liver-Blood Axis

**DOI:** 10.1101/2025.03.11.642626

**Authors:** Ran Ran, Merve Uslu, Mohd Farhan Siddiqui, Douglas K. Brubaker, Martin Trapecar

## Abstract

Understanding T cell clonal relationships and tissue-specific adaptations is crucial for deciphering human immune responses, particularly within the gut-liver axis. We performed paired single-cell RNA and T cell receptor sequencing on matched colon (epithelium, lamina propria), liver, and blood T cells from the same human donors. This approach tracked clones across sites and assessed microenvironmental impacts on T cell phenotype. While some clones were shared between blood and tissues, colonic intraepithelial lymphocytes (IELs) exhibited limited overlap with lamina propria T cells, suggesting a largely resident population. Furthermore, tissue-resident memory T cells (TRM) in the colon and liver displayed distinct transcriptional profiles. Notably, our analysis suggested that factors enriched in the liver microenvironment may influence the phenotype of colon lamina propria TRM. This integrated single-cell analysis maps T cell clonal distribution and adaptation across the gut-liver-blood axis, highlighting a potential liver role in shaping colonic immunity.

## Introduction

The gut-liver axis is an immunologically dynamic interface integrating signals from myriad microbial, dietary, and endogenous factors to maintain homeostasis and regulate responses to pathogens, commensal microbes, and foreign antigens. The colon alone harbors trillions of microbes which exert profound influences on host immunity, metabolism, and overall health^1,2^. The portal circulation is a critical component of this immunological dialogue, transporting nutrients and microbial metabolites from the colon to the liver, a major site of immune cell residence^3^. In parallel, lymphatic channels provide conduits for immune cell trafficking, establishing a complex network of interactions shaping local and systemic immunity^4^.

The gut and liver contain a large fraction of the body’s T lymphocytes^2,5,6^. Tissue-resident memory T cells integrate cues from the microbiota, dietary antigens, and inflammatory signals to balance tolerance and responsiveness^7^. Single-cell RNA sequencing (scRNA-seq) effectively maps T cell heterogeneity, providing resolution into transcriptional states, clonality, and tissue adaptations^8^. T cell receptor (TCR) clonality represents a critical dimension of adaptive immunity, reflecting how T cells expand in response to specific antigens and adapt to their environment. Profiling the single-cell TCR repertoire enables identification of clones undergoing selective expansion, potentially revealing immunodominant responses to commensals, dietary antigens, or pathogens^9,10^. These clonal signatures uncover how tissue-specific niches shape the T cell landscape and highlight differences between transient, circulating populations and stable, tissue-resident cells^11^. Coupling TCR clonality with transcriptional phenotypes enables pinpointing attributes that define the most successful clones.

Despite the immunological importance of these compartments, limited donor-paired data exists simultaneously characterizing the gut, liver, and circulating T cell populations. This critical gap confounds insights into T cells distribution, clonal expansion, and phenotypic specialization across interconnected tissues. Although prior studies provided critical frameworks for understanding tissue-specific immune dynamics and T cell subsets in the gut or liver individually^12,13^, heterogeneity in diet, lifestyle, genetics, and environmental exposures can significantly influence the composition and functional attributes of these immune cells^14^.

Emerging evidence of significant differences in transcriptional profiles, longevity, and responsiveness between tissue-resident and circulating T cells underscores the importance of understanding clonal relationships across the gut-liver-blood axis^15^. Tissue-resident T cells in the gut must discriminate between commensals and potential pathogens, while liver-resident T cells reside within a unique vascular and immunological microenvironment^13^. Previous scRNA-seq-based characterizations of gut or liver T cells underscored the complexity and variability of these immune populations, but have not resolved how these distinct niches interact within the same individual^8^.

Here, we characterized T cell populations from the large intestine, liver, and peripheral blood from the same donors, avoiding confounders factors of inter-individual variability to enable direct comparison of cell phenotypes, states, and clonality across tissues that have rarely, if ever, been profiled simultaneously. We integrated scRNA-seq and scTCR-seq to map transcriptional landscapes of T cells in each compartment and define their clonal architecture, thereby distinguishing tissue-resident subsets from circulating cells^16,17^. Unlike previous works, our strategy provides a uniquely robust framework to identify shared and divergent T cell signatures within the gut-liver axis of the same individuals.

## Result

### Site-Specific T Cell Phenotype Profiling by scRNA-Seq Along the Gut-Liver Axis

We performed scRNA-seq with paired TCR (αβ chain only) sequencing of CD3+ T cells from 4 locations (colon epithelium (intraepithelial lymphocytes, IEL), colon lamina propria (LP), liver (L), and peripheral blood (PB)) in 3 healthy human donors AJD3280, AJG2309, and AJKQ118. After quality control, we obtained 72,800 T cells with balanced integration from each location (Supp. Fig. 1a). Leiden clustering resulted in cell subsets that were manually annotated by their marker genes expression (Fig. 1a).

**Fig. 1.**
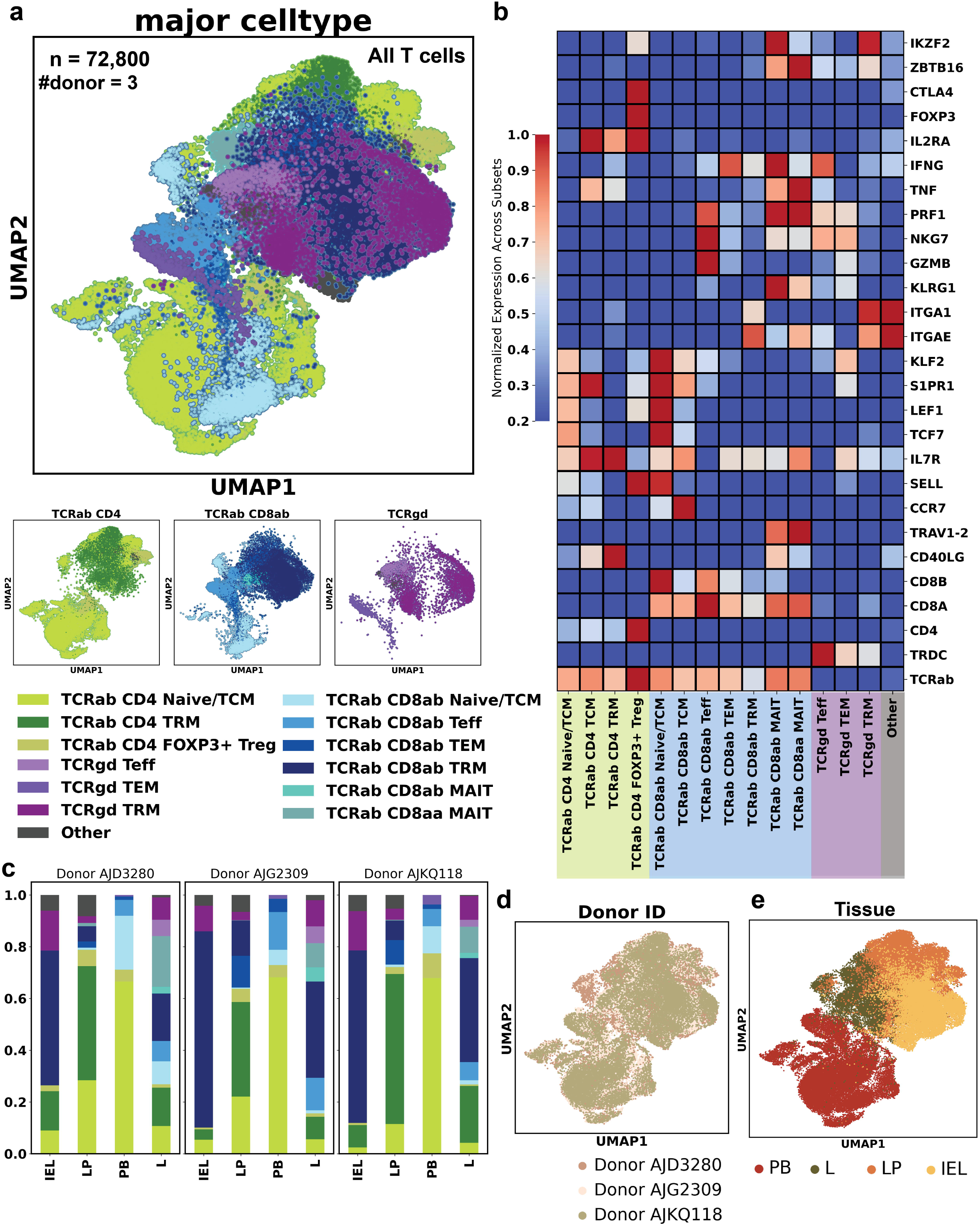

The dominant T cell subsets were consistent across the three donors, although some variations were observed (Fig. 1c, d and Supp. Fig. 1c). Specifically, TCRαβ CD8+ TRM were the most abundant T cell type in both the colonic epithelium and the liver, TCRαβ CD4+ TRM predominated in the colonic lamina propria, and TCRαβ CD4+ TCM were most prevalent in the blood. MAIT cells represented a larger proportion of T cells in the liver compared to the other three locations. γδ T cells are enriched in colon epithelium and liver, and only a few of them are found in the blood. Indeed, T cells isolated from PB have a distinct gene expression profile compared to the tissue-origin ones, which can be explained by their cell subset composition and its circulating nature (Fig. 1e). We calculated an activation module score from the expression of activation markers *IL2RA*, *CD38*, *ENTPD1*, cytotoxic molecules-encoding *GZMB*, *PRF1*, and proliferation marker *MKI67* (Supp. Fig. 1b). In blood, most T cells are at rest, whereas the intraepithelial and intrahepatic T cells, especially the TCRαβ CD8+ and γδ T cells, exhibited signs of activation even under homeostasis (Supp. Fig. 1b, e).

### Unique Signatures Define TRM at Different Sites

To identify site-specific gene expression signatures in TRM populations, we performed differential gene expression analysis comparing TRM cells across sites. Both CD4+ and CD8+ LP TRM cells displayed elevated expression of *ITGA1* compared to liver TRM (Fig. 2a-b), suggesting enhanced tissue retention in the colon. CD8+ LP TRM also uniquely upregulated *XCL1* and *XCL2* (Fig. 2a), chemokines critical for dendritic cell (DC) recruitment and cross-priming^18^, implying site-specific adaptations to local immune crosstalk. Interestingly, we saw CD4+ IEL TRM expressed higher levels of human leukocyte antigen-antigen D related (HLA-DR), part of the MHC class II determinants, compared to their lamina propria counterparts (Fig. 2a, Supp. Table 1). Gene set enrichment analysis (GSEA) suggested that LP TRM, relative to intraepithelial and liver TRM, exhibited increased interferon-α/β and chemokine-related signaling pathways (Adjusted P-value < 0.05; Fig. 2a, b, Supp. Table 2), consistent with their proximity to the cytokine-secreting antigen-presenting cells and the Gut-Associated Lymphoid Tissue (GALT).

**Fig. 2.**
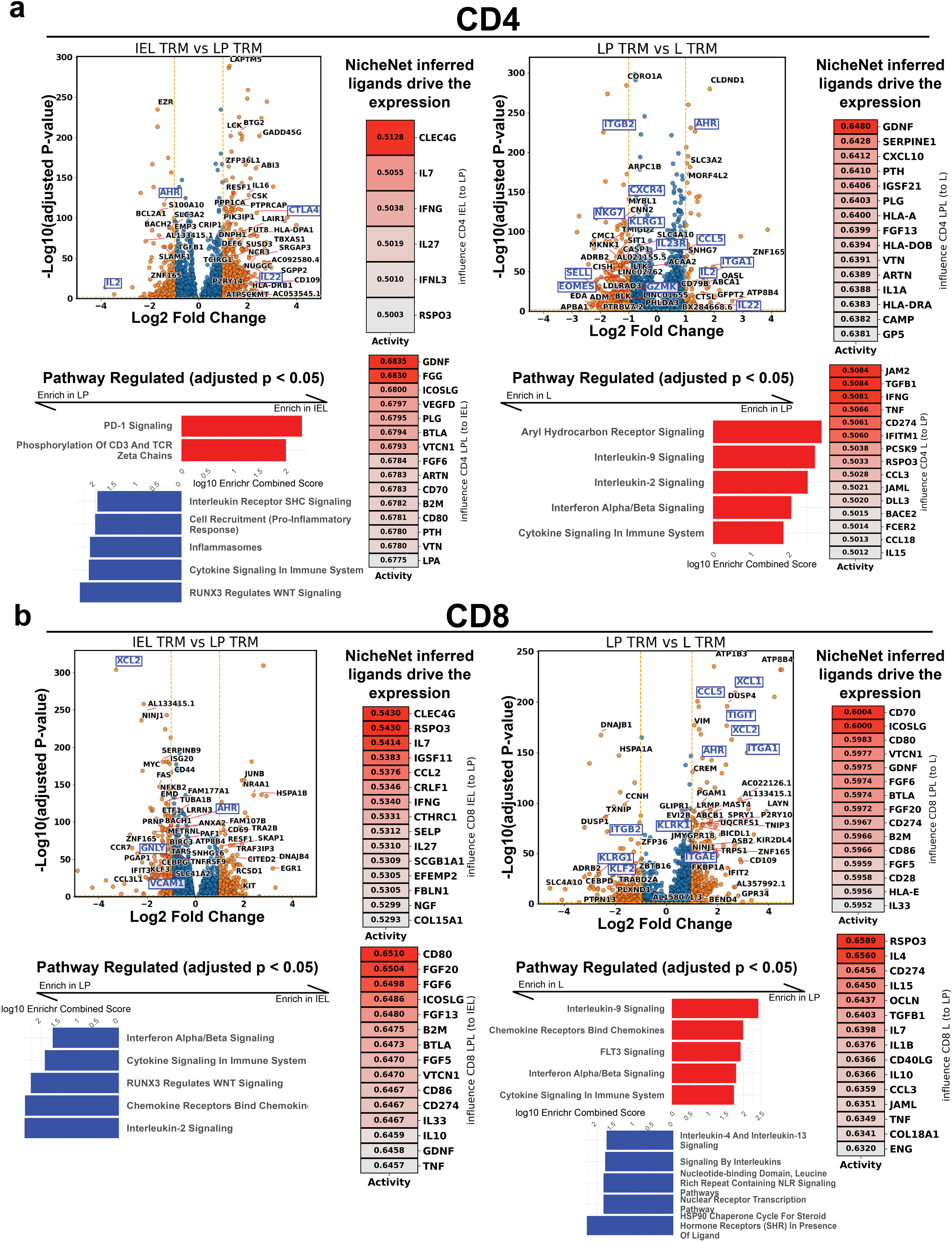

Since T cells interact with their environment through both direct cell-cell contact and secreted factors, we hypothesized that the observed differences in TRM gene expression across sites were driven, in part, by responses to ligands expressed by neighboring cells. To identify which neighbor cell types contribute to the site-specificity of the TRM, for each comparison (IEL vs LP, LP vs L), we input each site’s significantly upregulated genes into NicheNet^19^, which returned the predicted ligand regulatory power of causing the differential expression in the given set. NicheNet predicted ligands with lower confidence for IEL and CD4+ liver T cells with respect to LP T cells, but high confidence ligands in other cases (Fig. 2a, b, Supp. Table 3). The top ligands predicted to shape both the CD4+ and CD8+ TRM in lamina propria are related to extracellular matrix adhesion (ARTN, GDNF), interactions with the APC and/or B cells (CD80/86 provides co-stimulatory signal to CD28, CD70 binds CD27, ICOSLG binds ICOS on CD4+ T cells), and MHC-mediated activation implied by the B2M. For IEL TRMs, IL27, IL7, and IFNG are inferred to results in its unique expression. The liver-specific TRM features predicted to be influenced by a combination of interleukins (IL-4/IL-13, IL15) and cytokines including TNF, which promote hepatocyte proliferation and liver regeneration^20^.

Next, we explored which cell types in the environment may be the producers of these inferred phenotype-influencing ligands. We queried the cell type-averaged expression of our site-specific T cells and the non-T cells from two public annotated whole-tissue scRNA-seq datasets of human colon (epithelium and lamina propria)^17,21,22^ and liver^8^, and we used it to quantify the impact of local cell types on each TRM subset (Fig. 3, Supp. Fig. 2, Supp. Table 4). The CD8+ IEL T-influencing ligands are more expressed in epithelial cells, namely stem, goblet, and transit amplifying (TA) cells, indicating IEC-IEL connections. In the lamina propria, we determined that DCs contribute most to the CD8+ LP TRMs, whereas the fibroblasts and lymphoid stromal cells—cells near the GALT, like fibroblastic reticular cells—shaped the CD4+ LP TRMs together with other myeloid cells and stromal cells. Interestingly, liver sinusoidal endothelial cells (LSECs), a platform for adhesion of various liver-resident immune cell populations, were predicted to have the highest potential of resulting in the CD4+ liver TRM specialization, as described previously^23^, LSECs are the key liver immune regulators. On the other hand, in addition to the LSECs, the CD8+ TRMs in the liver are likely to be shaped by local macrophages^24,25^.

**Fig. 3.**
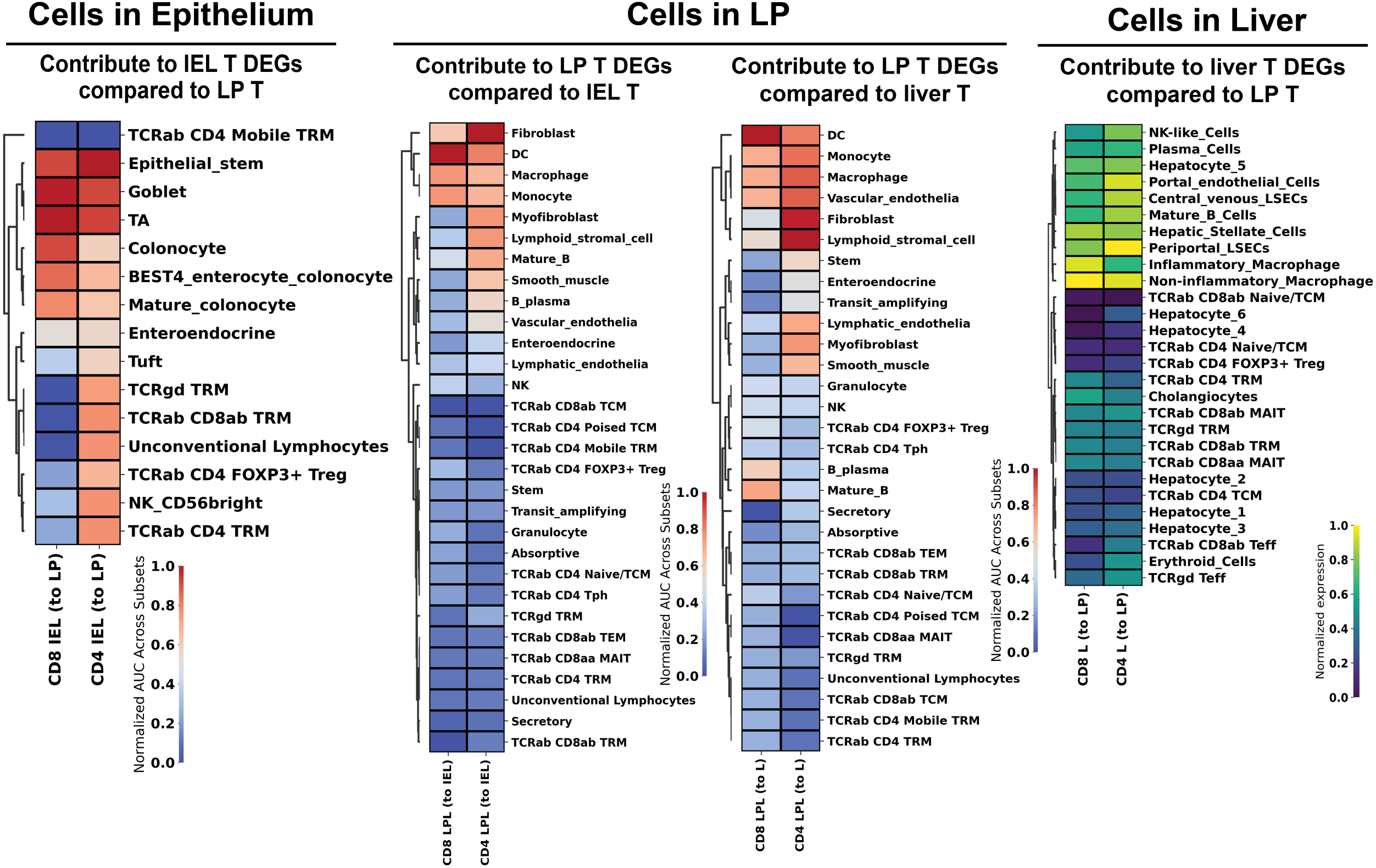

### Clonal Sharing Across Sites Reveals a Connection Between Colon- and Liver-Resident T Cell Populations

We investigated TRM population clonal relationships using paired TCRαβ-seq to determine whether the observed phenotypic differences reflected distinct clonal origins or the adaptation of shared clones to different tissue microenvironments. Assessing T cell residency, in part, depends on establishing these clonal relationships. In all donors, blood showed a distinctly higher diversity index of TCR repertoire, while T cells isolated from tissues were less diverse (Supp. Fig. 3a). Rare clones (clone with only 1 cell member) took up the majority (75-90%) of all 3 donors’ peripheral blood T cells repertoire, which explains the high diversity index of PB T cells (Supp. Fig. 3b).

We then compared “shared T cell clones” among sites, defined as clones present in multiple tissues. A maximum MOI of 0.32 indicates that the TCR repertoire at each site is predominately distinct (Fig. 4a). In all donors, IEL showed a moderate clonal connection with the LP. In Donor AJG2309, PB showed high clonal overlap with L, suggesting T cell migration from blood to liver. Interestingly, in two donors, LP also weakly overlapped with L, indicating marginal TCR repertoire similarity between colon and liver.

**Fig. 4.**
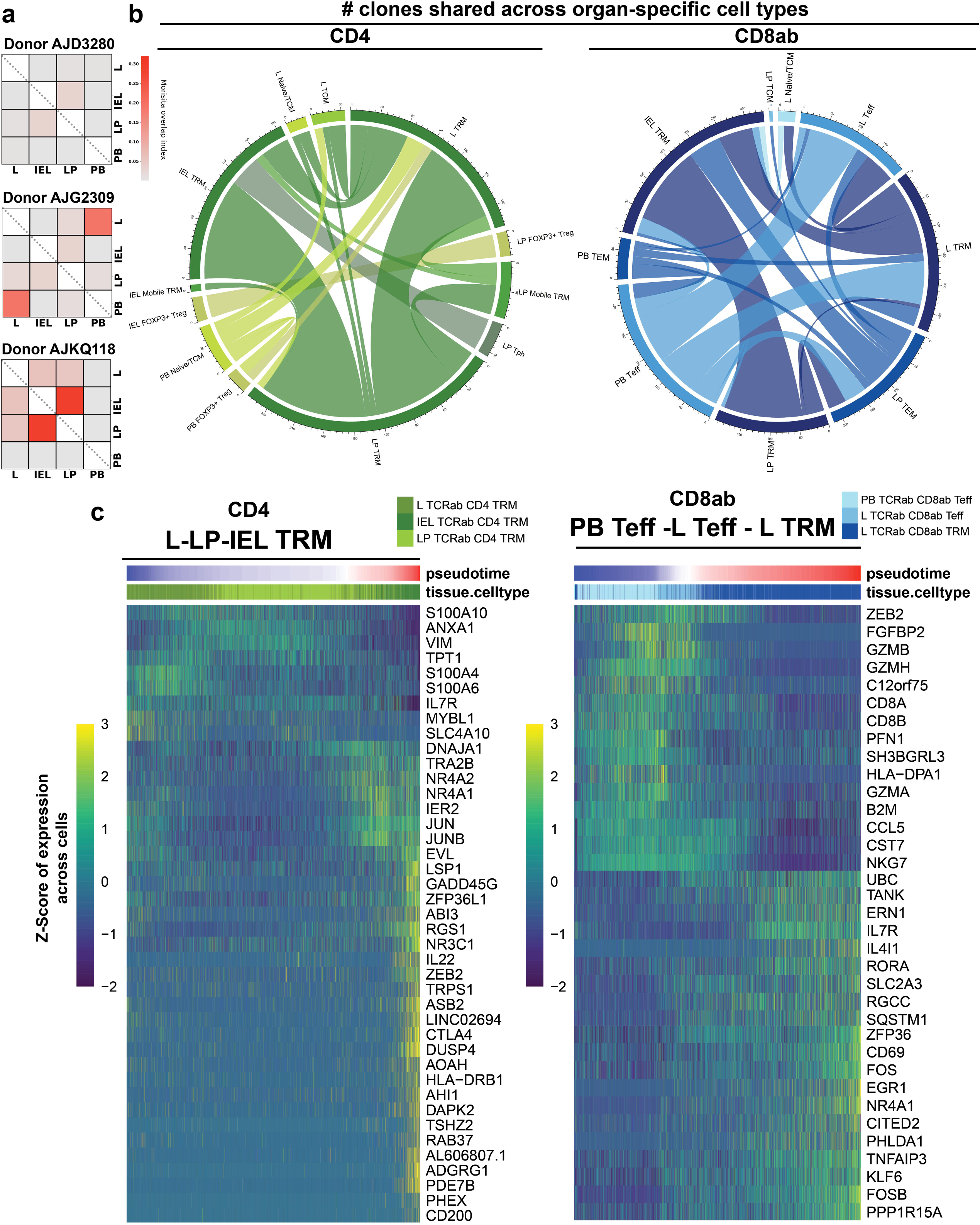

We increased the granularity of our analysis by focusing on the number of clones shared between site-specific cell subsets in all donors. Major clone sharing in CD4+ and in CD8+ T cells was observed among IEL TRM, LP TRM, and L TRM. Clones are also shared among PB Teff, L Teff, and L TRM (Fig. 4b). Interestingly, clones were shared between CD4+ IEL FOXP3+ Treg and CD4+ LP Treg, but not the circulating FOXP3+ Treg in the blood, indicating that the colon FOXP3+ Tregs are relatively isolated.

Clonal overlap between cell subsets suggests either parallel development or serial transitions. A transition among PB Teff, L Teff, and L TRM group is consistent with the theory of establishing TRM from effectors in the circulation^26–29^. While the TRM is believed to be tissue restricted^30^, the egress of TRM, especially under the pathological conditions, has been suggested in multiple studies given their presence in the blood or secondary lymphoid organ^30–34^. We explored transitions for the CD4+ TRM since these have been reported to exit the skin and present in the blood of healthy individuals^34^. We analyzed gene expression changes that might accompany the two hypothetical transitions using pseudotime analysis with Slingshot^35^. We inferred an increase in *CTLA4* and tolerance-related transcription factors^36,37^ *NR4A1* and *NR4A2*, along with a decrease in the calcium-binding protein-encoding genes *S100A4*, *S100A6*, *S100A10*, and the longevity marker *IL7R*, when transitioning from liver CD4+ T cells to intraepithelial CD4+ T cells (Fig. 4c). For CD8+ T cells, as they progress from PB Teff to L TRM, we observed a decrease in cytotoxic granzyme genes and *CCL5*, alongside an increase in *IL7R*, *RORA*, and *NR4A1* expression.

### Investigate The Most Expanded Clone at Each Location

To characterize clonal expansion, we ranked clones by frequency within each tissue and donor, grouping them into top 10, next 90 (11-100), and remaining (>100) clones, representing highly, moderately, and less expanded clones, respectively. In CD8+ cells, there is a higher proportion of cells from the top expanded groups at each location in each donor, meaning that CD8+ T cells are more clonally expanded than CD4+ T cells, in accordance with previous studies^12,38,39^ (Fig. 5a). For CD4+ T cells, most of the expansion happens at tissue, which is also generally the case in CD8+ except Donor AJG2309 who has an expansion of CD8+ cells in the blood. We determined the major phenotype of the most expanded cells at each location, which are CD8+ TRM in the epithelium, CD4+ TRM in the lamina propria (except Donor AJG2309, which is CD8+ TEM), and CD8+ Teff in the blood (Fig. 5b). In liver, CD8+ TRM were found to expand the most in Donor AJG2309 and Donor AJKQ118, in which the MAIT cells also contribute to a comparable portion of the top expanded cells. In fact, MAIT cells take up almost 80% of the most expanded cells in Donor AJD3280’s liver. Considering they were less than 20% abundance in the intrahepatic T cells (Fig. 1c), this distinctly high expansion rate indicated a MAIT-specific antigen-enriched environment in the liver.

**Fig. 5.**
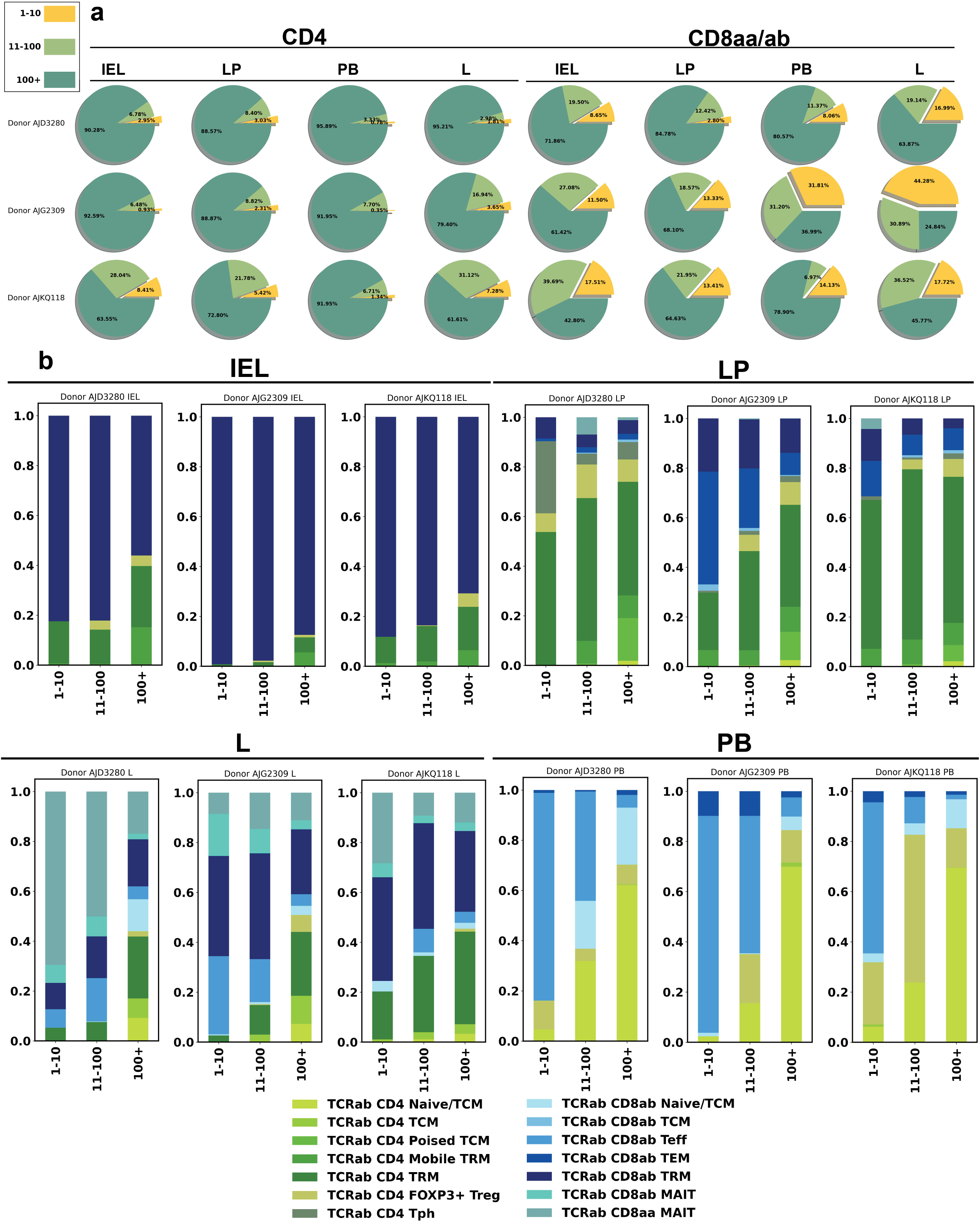

We compared the gene expression between cells in the most expanded clones (top 10) and least expanded clones (top 100+) of the same type at each location and donor. We found that *IL7R*, gene encodes the receptor of IL7 that promotes cell survival, is downregulated in the most expanded cells in Donor AJG2309’s liver CD8+ TRM, blood CD8ab Teff, and Donor AJKQ118’s blood CD8+ Teff (Fig. 6a, Supp. Fig. 4a, Supp. Table 5). This likely reflects short-lived effector differentiation along with the expansion after T cell activation because the effector-fate driving transcriptional factor T-bet^40^ (encoded by *TBX21*) inhibits IL7R^41,42^. We found *GPR183* higher in the least expanded clones in all cell subsets and statistically significant in Donor AJKQ118’s intraepithelial CD8+ TRM and liver CD8+ TRM. GPR183, an oxysterol receptor (EBI2) mediating immune cell migration^43–46^, was higher in the least expanded clones across all subsets, with significant differences in Donor AJKQ118’s intraepithelial and liver CD8+ TRM. We hypothesize that since expanded cells have been at least activated once, they downregulate this chemotactic receptor to avoid over-activation. As shown in Gianni Monaco et al.’s dataset, terminal effector memory CD8+ T cells indeed have significantly lower expression of *GPR183* than other T subsets^47^.

**Fig. 6.**
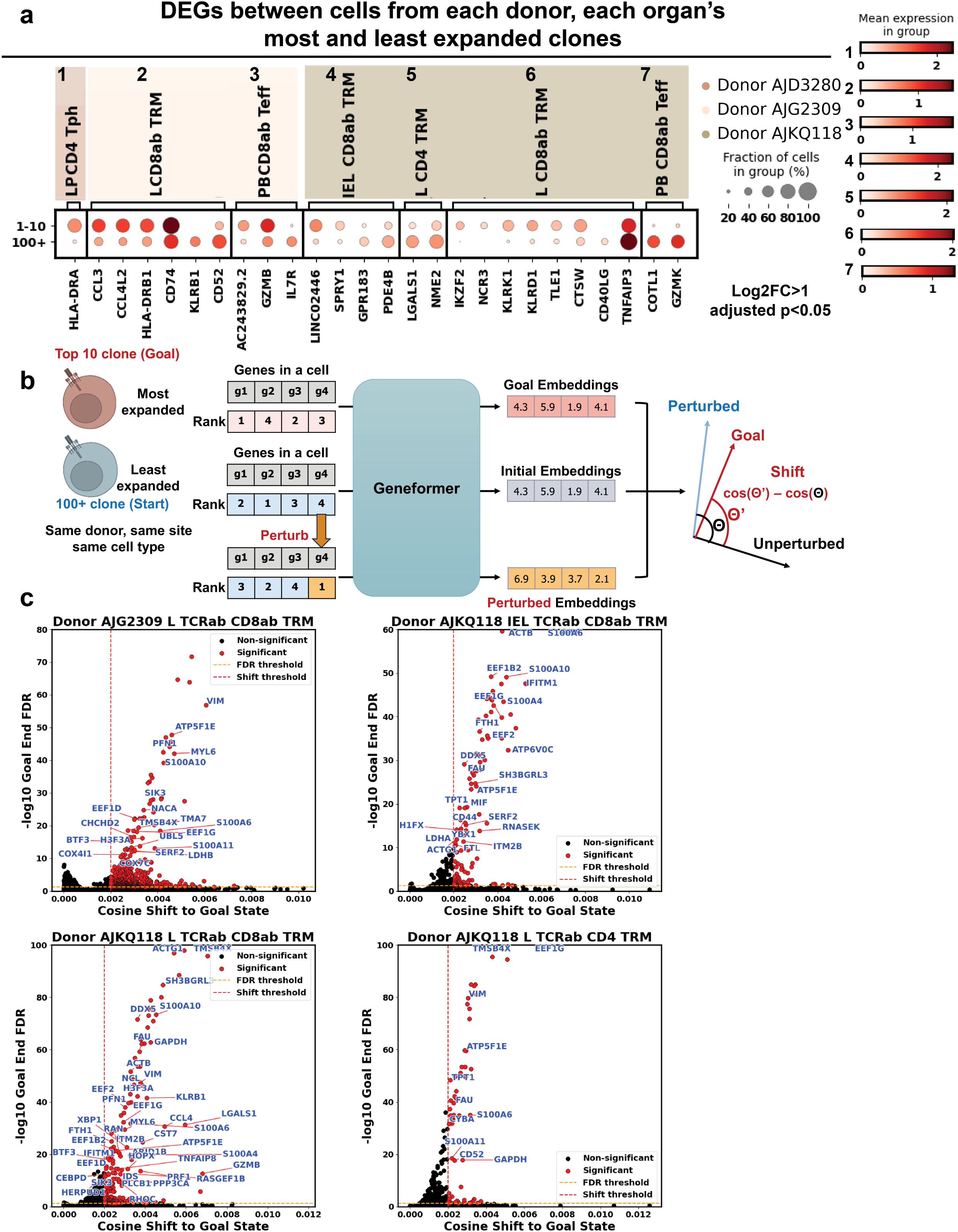

We examined whether the top 10 clones in each donor’s site were also present at other sites (Supp. Fig. 4b). The top clones of intraepithelial and liver T cells were confined locally. However, most expanded lamina propria and blood T cell clones—despite their low total frequency within each site, indicating relatively limited expansion—were often also found in the intraepithelial and liver compartments, respectively. This suggests an asymmetry in T cell migration and clonal expansion, with movement occurring primarily from the lamina propria to the epithelium and from the blood to the liver. We also analyzed *TRBV* gene usage preferences in T cells with varying degrees of clonal expansion across different sites (Supp. Fig. 4, 5, 6; Supp. Table 6). However, we found no consistent site-specific *TRBV* usage across all donors.

### In Silico Perturbation by Geneformer Predicts Potential Expansion Switch

Although expansion of TRM is mostly due the secondary exposure to the antigen of specificity, we asked whether this process can be assisted by modulation of gene expression. We applied Geneformer^48^, an in-silico perturbation tool developed from training large language model, to TRM subsets that were shown to have differential expression between the top expanded and least expanded clones to infer the molecular switch that may promote the expansion (Fig. 6b).

Most genes Geneformer inferred to be expansion-leading coordinate reprogramming of metabolic, cytoskeletal, and translational pathways (Fig. 6c, Supp. Table 7). For example, genes such as *ATP5F1E*, *COX4I1*, *COX7C*, and *LDHB* point to an increased reliance on mitochondrial respiration and glycolytic activity to meet the heightened energy demands of proliferating cells. Genes encoding cytoskeletal components like *VIM*, *MYL6*, and *PFN1*—along with regulatory kinases such as *SIK3*—indicate active remodeling of the cellular structure, which is important for cell division and mobility. The S100 family protein-encoding genes (*S100A6*, *S100A10*, and *S100A11*) modulate calcium signaling and other stress responses that further promote activation. Finally, genes involved in protein synthesis and regulation (*EEF1D, EEF1G, BTF3, TMA7, SERF2*, and *UBL5*) suggest increased translational support of cellular growth. Interestingly, some surface protein-encoding genes were detected in the list, including *KLRB1* in Donor AJKQ118 liver CD8+ TRM, which is a controversial co-stimulatory receptor with unknown mechanism that was reported to enhance T cell activation^49^. *CD52* in the same donor’s liver CD4+ TRM, which encodes a glycoprotein expressed on immune cell surface, was reported to provide a co-stimulatory signal^50^. We also found that overexpression of *CD44*—encodes an adhesion receptor that binds to hyaluronic acid in the extracellular matrix and a putative T cell activation marker—led the least expanded Donor AJKQ118 IEL CD8⁺ TRM population to a more highly expanded state *in silico*. This is reasonable, as CD44 binding to its ligand enables cell migration and provides costimulation to T cells^51–54^.

## Discussion

Our single-cell analysis of T cells across the human gut-liver-blood axis revealed significant differences between circulating and tissue-resident populations, both in their transcriptional profiles and TCR clonotypes. We found that the distinct microenvironments of the colon and liver imprint unique characteristics on their respective T cell populations.

A key feature of our study is the simultaneous isolation of T cells from the colon, liver, and peripheral blood of the same donors, enabling us to analyze clonal relationships across the gut-liver axis. The liver is a more blood-enriched organ than colon (∼100 ml/min per 100 grams of tissue^55^ compared to ∼35.4 ml/min per 100 grams of tissue^56^) where ∼75-80% of it is venous blood that portal vein collects from the intestine, stomach, spleen, pancreas, and gallbladder. This direct connection exposes liver-resident T cells to gut-derived soluble factors (metabolites, cytokines) and antigens. We found that clonal overlap between colon and liver T cells, while present under homeostatic conditions, varied significantly between individuals. This inter-individual variability may reflect differences in prior exposures, disease history, genetics, or other factors that influence clonal expansion and the establishment of tissue-resident niches. Nevertheless, because of the rich blood flow, most expanded blood T cell clones were found in the liver, although the absolute cell count was relatively low.

Our study extends previous work on colon-resident T cells by separately analyzing intraepithelial lymphocytes (IELs) and lamina propria T cells, providing single-cell resolution gene expression profiles for these distinct compartments. While much of the existing literature on IELs focuses on mouse models or the human small intestine, our work specifically profiles human colonic IELs. Murine studies yield an estimated αβ/γδ IEL T cells ratio about 2:1 or even 1:1 in small intestine and 5:1 in colon^57,58^. Our study reported a much smaller percentage (< 20%) of gamma delta IEL T cells in the human colon, which agrees with previous human colon IEL studies^59–61^.

Compared to CD4+ TRM cells in the lamina propria, CD4+ IEL T cells expressed higher levels of *HLA-DR*, which encodes MHC class II molecules. While classically associated with antigen presentation by professional APCs, *HLA-DR* expression is also found on activated CD4+ T cells^62–64^, and is a marker of Treg subsets with distinct functions^64–67^. It is unclear whether *HLA-DR* expression on CD4+ IEL TRM confers suppressive capabilities like Tregs. One study showed that blocking HLA-DR did not impair contact-dependent suppression by certain T cell subsets or the interaction between HLA-DR and its ligands CD4+ or LAG-3 in maintaining local homeostasis^64^. It is intriguing to ask whether these MHC II molecules expressed on IEL T cells can present antigens. In the epithelium, it is thought that the epithelial cells serve as non-conventional APCs that present antigen to the intraepithelial T cells^68–71^. Expression of MHC II is seen in the epithelial cells collected by Gut Cell Survey (Supp. Fig. 2). Since MHC class II on CD4+ T cells presents antigens in humanized mice^63^, we hypothesize that CD4+ IEL T cells might present antigen to *each other* within the epithelial compartment, potentially contributing to local immune regulation.

During the late 90s, several studies used PCR to amplify the rearranged TCR region to explore the IEL TCR repertoire^72–76^. They defined a "clone" as T cells whose PCR-amplified rearranged TCR gene was detected as an identical product. They reported that both the human small intestine and colon IEL are largely oligoclonal, with the epithelium compartment favoring a few specific TRBV gene. However, the old method of clonotyping is greatly limited: antibody approaches target TCRβ and TCRα variable regions cannot differentiate clones with identical V regions, needless to say the technical error associated with amplification at the time. In our work using scTCR-seq, though several IEL clones exhibited significantly more expansion than others, none of the CD4+ and CD8+ αβ IEL T cells exhibit distinctly strong oligoclonality compared to the other sites. A study reported multiple human colon segments TRBV sequences Simpson’s D diversity index averaged between 0.01 and 0.1, which is considered highly diverse^77^, and similar results that supports the polyclonality was also observed in human infants and children’s SI and colon^78^.

Regarding T cell migration between the epithelium and lamina propria, live animal imaging reports frequent leaving and entering of γδ intraepithelial T cells^79^. Tamoxifen-inducible live cell tracking in murine small intestine showed γδ T cells and FOXP3+ Treg migrating between lamina propria and epithelium over 24 hours, showing a steady exchange^71^. It was reported that when small intestinal epithelium was denuded, LP T cells moved across the pores on the basal membrane, although their ability to pass through an intact epithelium is low^80^. Based on the anatomical proximity of IELs and LP T cells, and the fact that blood vessels supplying the intestine are located within the lamina propria, we expected to observe some degree of clonal overlap between αβ IELs and αβ LP T cells. Indeed, previous studies in both mice and humans have reported shared clones between these compartments^72,81^. However, if frequent bidirectional exchange occurred, we would anticipate a much higher degree of clonal overlap. Our analysis, in contrast, revealed relatively limited clonal sharing between αβ colon IELs and LP T cells, particularly among the most expanded clones. As in previous mouse studies of small intestinal γδ IEL^81^, our analysis supports that αβ colon IEL T cells, especially those from the most expanded clones, are relatively confined to the epithelium, although marginal translocation exists and mainly exists directionally from lamina propria to epithelium given the most expanded LP clones can be found in epithelium but seldomly the other way around.

The limited clonal exchange between IELs and LP T cells, particularly the restricted movement from epithelium to lamina propria, likely contributes to the observed differences in cellular composition across the epithelial basement membrane. The epithelium provides a unique microenvironment, where IELs interact directly with epithelial cells^79,82–84^ and are exposed to luminal microbes and antigens^81^. These interactions influence IEL mobility, activation status, and gene expression profiles. Our ligand inference using NicheNet predicts epithelial tight junction protein occludin (OCLN) to be one of the ligands that regulate IEL T cell expression, which was proved to be vital for the migration of γδ IEL T cells^79^. Intriguingly, our analysis also indicates that ligands uniquely enriched in liver, such as fibrinogen gamma chain (FGG), vitronectin (VTN), lipoprotein A (LPA), and plasminogen (PLG), may affect the phenotype of T cells in the lamina propria. After querying NicheNet’s ligand-receptor interaction library, these 4 ligands are defined to bind to the integrins, or in our case, the ITGB1 and ITGB2 whose RNA is expressed by the LP TRMs. We hypothesize that T cells may encounter these liver-enriched ligands within the hepatic microenvironment and subsequently migrate to the colonic lamina propria, where integrin signaling further contributes to their tissue-resident phenotype and function^85,86^. This suggests a potential mechanism by which the liver can indirectly influence the characteristics of colon-resident T cells.

Our findings reveal a previously unappreciated connection between the liver and colonic immune landscape, suggesting that systemic factors, potentially originating in the liver, can shape the phenotype and function of gut-resident T cells. Future studies should explore the precise molecular mechanisms underlying this inter-organ communication and its potential role in both health and disease, including inflammatory conditions beyond the gut-liver axis.

## Method

### Human Subjects

Human colon tissue, liver tissue, and blood samples were purchased from LifeNet Health LifeSciences. The first non-diseased donor (AJD3280) was a Caucasian female aged 28 with an BMI of 20. The second non-diseased donor (AJG2309) was a Caucasian male aged 43 with an BMI of 24.3. The third non-diseased donor (AJKQ118) was a Caucasian male aged 56 with an BMI of 19. The tissue was transported in the University of Wisconsin perfusion media and blood in purple top tubes on ice. It arrived in our facilities in under 24 hours of cold ischemia time and under 6 hours of warm ischemia time. Samples were processed immediately upon arrival.

### Processing And Immune Cell Isolation from Colonic Tissue

Tissue was processed based on a previously established protocol^87^. In brief, colon tissue was washed with pre-rinse buffer containing Hank’s 1X Balanced Salt Solutions (HBSS) without calcium and magnesium (Cytiva, Cat no: SH30588.01) and penicillin-streptomycin-amphotericin B (PSA) (Lonza, Cat no: 17-745E). First, adipose tissue was removed, and then the epithelium was peeled off from the muscularis externa with scissors. Each trimmed colon tissue was cut into small pieces with scissors and placed into small tissue culture dishes (21.2 cm^2^) (Olympus Cat No: 25-260). Tissues were minced into fine pieces with an approximate size of 5mm using the McIlwain Tissue Chopper (Cavey Laboratory Engineering Co. Ltd). On the day of isolation, we prepared i) the rinse buffer containing RPMI-1640 (Gibco, Cat no: 11875-093), 5% HI-FBS (Cytiva HyClone, Cat no: SH30910.03) and 1% Penicillin-Streptomycin (PS) (Gibco, Cat no: 15-140-122), ii) pre-digestion buffer with 5mM EDTA ((Invitrogen; Cat no: 15575-038), 1 mM DTT (Teknova; Cat no: D9750), 10 mM HEPES (Gibco, Cat no: 15630-080), 1% PSA (Lonza, Cat no: 17-745E), 0.1 % BSA (Fisher, Cat no: BP1600-100) in 1x HBSS without calcium and magnesium (Cytiva, Cat no: SH30588.01), iii) the digestion buffer was prepared as follows: 5 mg/ml Collagenase D (Roche; Cat no: 11088858001), 0.5U/ml Dispase (Gibco; Cat no: 17105-041), 30 μg/ml DNAse I (StemCell Technologies, Cat no: 07470), 10 mM HEPES (Gibco; Cat no: 15630-080), 1% PSA (Lonza, Cat no: 17-745E) in HBSS with calcium and magnesium (Cytiva, Cat no: SH30030.01). All buffers were warmed at 37°C.

Tissues were resuspended with 15 ml of prewarmed predigesting buffer by inversion. Then, it was incubated horizontally for 20 min at 37°C under continuous rotation (140 RPM) using an incubating shaker (Thermo Scientific, MaxQ 420HP). After that, the tissue pieces waited to settle to the bottom of the 50-ml tube, and then the supernatant containing the stripped epithelium was transferred into a new 50-ml conical tube placed on ice using a pre-wetted 25ml serological pipette. Another 15ml of fresh prewarmed predigesting buffer was added to the colon tissue pieces and incubated horizontally for 20 min at 37°C under continuous rotation (140 RPM) using an incubating shaker (Thermo Scientific, MaxQ 420HP). After the second round of epithelial stripping, the tube was mixed gently for 10 seconds using a vortex and then waited to be settled at the bottom of the 50-ml conical tube. The second supernatant containing the stripped epithelium was collected and combined with the first round into a 50ml conical tube placed on ice using a pre-wetted 25 ml serological pipette. The 100μm cell strainer was washed with 5-10ml of rinse buffer, and the combined supernatant was passed through a 100μm cell strainer placed on a new 50-ml conical tube. 15 ml of ice-cold Rinse buffer was added to the tissue pellet, then centrifuged at 400xG for 10 minutes. The supernatant was transferred to a tube with a 70μm cell strainer and combined with previous supernatants. This combined cell suspension containing stripped epithelium and intraepithelial lymphocytes was centrifuged at 500g for 10 minutes. The pellet was resuspended with rinse buffer. Total cell number and cell viability were determined with an automated cell counter (Denovix, CellDrop FL) using trypan blue staining.

The remaining tissue pieces were transferred to gentleMACS^TM^ C tubes (Milteny Biotec, Cat no: 130-093-237) and washed with rinse buffer. Then, they were centrifuged at 500G for 5 min. 2.5 ml of prewarmed digestion buffer was added to tubes and incubated for 30 min at 37°C under continuous rotation (140 G) using an incubating shaker. Then, mix for 10 using a vortexer. The C tubes with the digested pieces were placed into the gentleMACS^TM^ Dissociator, and the program ‘‘m_intestine_01’’ was used twice. The tubes were briefly spun, and samples were aspirated using a 5-ml syringe and a blunt 20G needle three times. The cell suspension was passed through a 70μm cell strainer and washed with rinse buffer. The same digestion step was repeated by adding another 2.5 ml pre-warmed digestion buffer to the remaining tissue pieces and incubating for 30 min at 37°C under continuous rotation (140 G) using an incubating shaker. Then, mix for 10 using a vortexer. The C tubes with the digested pieces were placed into the gentleMACS^TM^ Dissociator, and the program ‘‘m_intestine_01’’ was used twice. The tubes were briefly spun, and samples were aspirated using a 5-ml syringe and a blunt 20G needle three times. The cell suspension was passed through a 70μm cell strainer and washed with rinse buffer. All collected lamina propria cells were combined, and total cell number/viability was determined with an automated cell counter (Denovix, CellDrop FL) using trypan blue staining.

The IELs and LP cell suspensions were centrifuged at 500G for 15 min separately. The cell pellets were resuspended in a DNAse solution prepared with 100 μg/ml DNAse I (StemCell Technologies, Cat no: 07470) in RPMI-1640 and incubated at room temperature (RT) for 15 min. An equal volume of rinse buffer was added, and the suspensions were centrifuged at 300G for 10 min.

### Isolation Of Liver-Tissue Resident Immune Cells from Liver Tissue

The liver tissue was washed with RPMI buffer to remove excess blood upon receiving the liver tissue. The liver wedge was cut into fine pieces of approximately 3-5mm using a scalpel and sharp scissors at room temperature. Before the day of isolating the immune cell from liver tissue, different buffers were prepared: i) rinse buffer (RPMI-1640+ 1% PSA+ 5% FBS), ii) Digestion buffer (Collagenase D 0.5mg/ml) + DNase 30ug/ml + HEPES 10mM + PSA 1% + HBSS (Ca+ Mg+)) and iii) EGTA buffer (10mM EGTA+ HBSS (Ca+, Mg+)) and stored them in 4C. The next day, all the buffers were placed in a 37°C water bath. Tissues were transferred to 30 ml of warm RPMI-1640 media and centrifuged for 10 minutes at 300G room temperature (RT). The supernatant was discarded, and the pellet was resuspended into the 20 ml EGTA buffer. The sample was shaken for 20 minutes at 37C in the shaker incubator. The samples were added with 20ml of Rinse buffer to make the 40ml total volume. The sample was centrifuged for 10 minutes at 300g RT. The pellet was resuspended in a 2.5ml digestion buffer and shaken for 20 minutes at 37°C in the shaker incubator. Thereafter, digested samples were transferred to Miltenyi C-tubes to homogenize the tissue using GentleMACS tissue dissociator preset program E.01. The homogenized samples were transferred to the 50ml tube using a 100μM pore size filter. We rinsed the filter with rinse buffer and made the 40ml total volume. The samples were then centrifuged for 10 minutes at 800G at 4°C. The supernatant was decanted, the pellet was resuspended in 30 ml 33% Percoll solution (RPMI67% 33% Percoll) (Cytiva, Cat no: 17544501), and the samples were spun at 1000g for 20 minutes at RT with acceleration and brake set to 1 and 0, respectively. The supernatant was gently decanted, and the pellet was resuspended in 5ml AKC lysis buffer (Gibco, Cat no: A10492-01) for 1 min and then centrifuged 800g for 5 min at 4C. The supernatant was gently discarded, and the pellet was resuspended in RPMI solution (RPMI+ 1% PSA+ 10% FBS+ Glutamax) and again centrifuged at 800g for 5 minutes at 4°C. The pellet was resuspended in DNASE 1 solution, incubated for 15 minutes at RT, and centrifuged at 500g for 10 mins at RT. The supernatant was discarded, and the pellet was resuspended in 1-2ml (based on the density of the pellet) PBS solution. Total cell number/viability was determined with an automated cell counter (Denovix, CellDrop FL) using trypan blue staining.

### Cell Sorting

Gut IELs and LP cells, PBMCs and liver immune cells were stained with LIVE/DEAD fixable green, fluorescent reactive dye (ThermoFisher, Cat no: L34970) according to the manufacturer’s protocol. In the meantime, an antibody mixture was prepared in BD Horizon Brilliant Stain Buffer ((BD Bioscience, Cat no: 563794). Then, cells were further stained with BV510-conjugated mouse anti-human CD45 (1:20) (BD Bioscience, Cat no: 563204) and PE-conjugated mouse anti-human CD3 (1:10) (BD Bioscience, Cat no: 555333) followed by incubation with human BD Fc block (1:100) (BD Bioscience, Cat no: 564220). After staining at 4°C for 20 min, cells were washed with FACs buffer containing 1% BSA in PBS and centrifuged at 300G for 10 min at 4°C. Cells were resuspended in FACs buffer, and CD45+ CD3+ cells from IELs, LPs, PBMCs and Liver-immune cells were separately sorted by fluoresce-activated cell sorter (Cytoflex SRT, Beckman Coulter).

### Single-cell RNA Sequencing

The Chromium Next GEM Single Cell 5′ Reagent Kit v2 (10x Genomics) was used for scRNA-seq. Sorted CD45⁺CD3⁺ T cells from the gut, liver, and blood were loaded into separate lanes of the Next GEM Chromium Controller (10x Genomics) for encapsulation. Single-cell libraries were generated according to the manufacturer’s protocols. TCR sequencing libraries for TCRαβ were prepared using the V(D)J Enrichment Kit (10x Genomics), following the manufacturer’s instructions. Library sequencing was performed at the Johns Hopkins SOM Single Cell and Transcriptomics Core.

### Data Quality Control and Pre-Processing

Seurat^88^ and Scanpy^89^ were used for downstream scRNA-seq data analysis. For quality control, low-quality cells were dropped based on their extremely low UMI counts (<1300/600/700 for intraepithelial T cells compartment, <700/900/900 for lamina propria T cells, <700/1200/600 for liver T cells, and <9090/1300/1100 for T cells isolated from peripheral blood. Reported in the format of donor AJD3280, AJG2309, and AJKQ118), high mitochondrial gene counts (>20%), and a small number of uniquely expressed genes (<200) to filter out empty droplet and dead cells. Python package doubletdetection^90^ was used to remove droplets. Scanpy function score_genes_cell_cycle was used to assign each quality-controlled cell with a cell cycle score based on the expression of cell cycle phase genes defined by Triosh et al^91^. Raw counts of quality-controlled cells, along with their cell cycle score and mitochondrial fraction, were input into R for normalization and donor integration. We performed SCTransform for each donor, each organ individually and then integrated different donors together. Gene raw counts were fitted into negative binomial distributions whose expectation was a function of the total count of cells. The coefficients of gene’s general linear model are further regularized by the kernel regression with the coefficient of other genes that have similar average expression. For our study, cell’s G2/M phase score, S phase score, and mitochondrial percentage were also included in the regression formula to remove the confounding effect of these variables on the gene expression. Regressed gene expression calculated from the regularized generalized linear model coefficients was referred as “SCTransform-corrected counts,” and log-transformed SCTransform-corrected counts were used for later visualization. The Pearson residual of a cell’s observed gene expression to the SCTransform-corrected counts was used for integration and later performing principal component analysis (PCA, 50 pcs kept). Data integration was done by using Seurat. The top 3000 genes that were the most variable across all donors’ organs’ data were selected to define the cell-anchors-finding space and then perform the integration based on canonical correlation analysis. Pearson residuals calculated by the SCTransform were corrected in according to the cell anchors. After PCA, Leiden clustering was performed based on the computed neighborhood graphs (50 pcs, size of neighborhood equals 15 cells) to reveal the general subtypes. The initial clustering resolution is set at 0.8. In order to delineate the T cell subtypes, further subclustering was performed on each subcluster at a resolution range from 0 to 1 (detailed resolution for each step of subclustering was recorded in the published code). UMAP was performed on the neighborhood graph to visualize the clustering (parameters set to scanpy default). These analyses—integration, PCA, UMAP, and cell type annotation—were performed separately for each site (with donor-integrated data). For the ultimate visualization, the four site-level datasets were re-integrated using SCTransform and Seurat, followed by PCA and UMAP, to generate a combined visualization while preserving the site-specific variation.

We classified cells as either TCRαβ or TCRγδ based on the detection of αβTCR transcripts and the expression of the γδ T cell marker *TRDC* (sufficient when CD3+ T cells are guaranteed)^92^ (Fig. 1b). TCRαβ cells were delineated into CD4+, CD8αα+, and CD8αβ+ subsets based on the expression of *CD4*, *CD8A*, and *CD8B*. We defined six major T cell subtypes as: (1) Naïve/Central memory T cells (Naïve/TCM) expressing *SELL*, *CCR7*, *TCF7*, *IL7R*, *SP1R1*, and no tissue residencies markers such as integrins *ITGAE* and *ITGA1*, showing their lymph node homing ability, the self-renewal and differentiation potential, long lives, and circulating ability. (2) Effector memory cells (TEM), which feature impaired *SELL* and *CCR7*, have effector marker *KLRG1*, have *S1PR1* or its transcription factor-encoding gene *KLF2* implies the potential to circulate in the blood, and *IL7R* indicates their long life. (3) Tissue-Resident Memory T cells (TRM), expressing high levels of *IL7R*, *ITGAE*, and *ITGA1*, and low levels of molecules associated with tissue egress, such as *S1PR1* and *CCR7*. As previously reported, CD4+ TRM expressed lower levels of CD49a (ITGA1) compared to CD8+ TRM^93^. (4) Effector T cells (Teff), characterized by low *IL7R* expression (indicating a short-lived phenotype), high *KLRG1* expression, and cytokine production. (5) FOXP3+ regulatory T cells (FOXP3+ Treg) in the CD4+ class featuring high *IL2RA*, *CTLA4*, and *FOXP3* expression. (6) Mucosal-associated invariant T cells (MAIT), which have a unique semi-invariant TCR α chain that uses *TRAV1-2*. We also identified relatively rare cell types such as CD4+ T peripheral helper cells expressing low *CXCR5* but high *CXCL13*^94^ in lamina propria and KIR^+^ γδ T that is the mouse Ly49^+^ Treg-equivalent in liver^95^ (Supp. Fig. 1b). Cells that did not fit into these defined categories were classified as unconventional lymphocytes. Activation score was assigned based on the aforementioned activation-related genes using scanpy function score_genes. For cell subsets that more than 30% of the cells have a positive activation score, we labeled them as activated. For those Naïve/TCM-agreeing sub-clusters labeled as activated, we reannotated them as TCM to avoid analyzing them with the supposedly quiescent naïve T cells.

### T Cell Receptor Information Pairing and Clone Assignment

TCR sequences were mapped to the scRNA-seq data based on each cell’s identifier. Only cells with at least one pair of productive TCR α- and β-chains captured in the TCR-seq were included in the TCR analysis, which yielded 26,297 qualified T cells from all three donors. For cells with more than one α- or β-chain sequenced, the chain with the higher consensus count was selected for downstream clonotype assignment. Cells sharing the same TRAV, TRBV, CDR3α, and CDR3β within the same donor were considered part of the same clone. Despite the exclusion of some cells from clonotype analysis, overall cell type composition remained comparable to the full dataset (Supplementary Fig. 1d). For each donor’s unique site, the top expanded clone was defined as the clone with the highest number of cells at that site. The Chao1 diversity index was estimated for the TCR repertoire in each donor and each site using the R package immunarch^96^. The rare clonal proportion was visualized using the repClonality function in immunarch. The Morisita Index between two sites within a donor was calculated based on the number of cells at each site belonging to clones present at both sites, relative to the total number of cells in each site. Chord plots were generated using the R package circlize, where links between site-specific cell subtypes in a donor represent the number of shared clones. Links corresponding to fewer than six clones were omitted for clarity. Overlaps between the top 10 clones at each donor’s site and clones from other sites within the same donor were visualized using the trackClonotypes function in immunarch.

### Differentially Expressed Gene Analysis

Differentially expressed gene (DEG) analysis was performed among TRM subsets across sites and between cells from the most expanded clones and least expanded clones within the site using Monocle 3^97^. This involved fitting a generalized linear model (GLM) to SCTransform-corrected counts, assuming each gene’s expression in each cell follows a quasi-Poisson distribution, with its mean and variance as functions of cell identity (site for TRM comparison, or whether the cell belongs to the most expanded clone for the most/least expanded comparison).

Log_2_ fold change was calculated from the log_2_-transformed ratio between the coefficient of the cell identity variable and the intercept (baseline). The *p*-value was computed using the Wald test on the coefficient to determine whether it was significantly different from zero. Benjamini-Hochberg correction was applied for multiple comparison corrections. The exact calculation procedure is described in the Monocle 3 documentation and our previous work^98^. Genes with adjusted *p*-values < 0.05 and an absolute log_2_ fold change > 1 were considered differentially expressed. Genes expressed in fewer than 10 TRM cells across all donors and sites were excluded from the DEG analysis.

Donor-site-specific DEG analysis between the most expanded and least expanded cells of the same cell subtype was performed using the Wilcoxon rank-sum test, a robust non-parametric test implemented as rank_genes_groups in scanpy for efficiency considerations. This analysis was applied to cell types that contained at least 10 cells from the most expanded clones and had a most expanded/least expanded ratio not greater than 10, ensuring a meaningful rank-sum test. Genes with adjusted *p*-values < 0.05 and an absolute log_2_ fold change > 1 were considered differentially expressed.

### Gene Set Enrichment Analysis

We performed gene set enrichment analysis (GSEA) using the **R** package enrichR^99^, with Reactome 2022 as the ontology database. The enrichment calculation method used by enrichR is detailed in the original author’s work. Briefly, enrichment *p*-values are computed using Fisher’s exact test, followed by multiple comparison correction using the Benjamini-Hochberg method. Differentially expressed genes (DEGs) from our analysis were input into the enrichr function for term enrichment calculation. Terms with adjusted *p*-values < 0.05 were considered significant. We visualized a subset of significant terms related to immunology, while the full list of enriched terms is provided in Supplementary Table 2.

### NicheNet Ligand Inference and Cell Type Ligand Enrichment Calculations

NicheNet^19^ (v2.2.0) was applied to infer the ligands most likely responsible for differential gene upregulation in each compartment across DEG comparisons. DEGs with adjusted p-values < 0.05 and log_2_ fold changes greater than 0.5 were used as input for NicheNet. The integrated ligand-receptor pairs, signaling network, and gene regulatory networks (prefix 21122021) were obtained from the package’s repository. Genes expressed in at least 1% of the cells within TRM subsets were considered as background. Potential ligands were defined as ligands of NicheNet-documented receptors whose encoding genes were expressed in at least 1% of the upregulated cell subsets. The top 15 ligands with an AUROC > 0.5 were reported in the main figure, while the top 30 were used for quantifying their enrichment in neighboring cell types. If fewer than 15 ligands met the AUROC > 0.5 threshold, all qualifying ligands were visualized and included in the enrichment analysis. The full list of ligands is available in Supplementary Table 3. Gut and liver profiles were obtained from the Gut Cell Survey Pan-GI Cell Atlas^17,21,22^ and GSE115469^8^ datasets. These datasets had already been pre-processed and normalized by their providers. AUC calculations were performed after averaging gene expression by cell type. A gene was included in the averaged profile only if it was expressed in at least 1% (for public reference data, which has a large sample size) or 10% (for T cells in our data) of a given cell type. Cells that were likely misannotated by the data providers (e.g., oral mucosa fibroblasts in the large intestine, fibroblasts in the epithelium, etc.) were removed (details provided in the script). Additionally, cell types with fewer than 30 cells were excluded due to their low statistical power. For AUC calculations, non-T cells from the public datasets and T cells from our study were concatenated, as AUC calculations were conducted for each cell type individually. For each cell type, we iterated down the ranked gene list (ranked by their average expression in the given cell type, descending order) to recover NicheNet-inferred ligands, stopping when encountering a zero-expression gene. The final area under the curve was computed using the auc function from sklearn.metrics.

Admittedly, small molecules produced by epithelial cells may cross the basement membrane into the lamina propria. Therefore, we considered non-T cells from both the lamina propria and the epithelium when identifying ligand-producing cells that could potentially regulate LP T cells. However, conventional epithelial cells generally do not perform robust reverse transcytosis from the lamina propria back into the epithelial layer in a way that would efficiently target IEL T^100^ Additionally, the basement membrane itself may block the diffusion of cytokines and growth factors, as has been suggested in tumor studies^101^. Thus, we only considered the influence of IEL cells and epithelial cells on IEL T cells.

### Pseudotime analysis

Pseudotime analysis was performed using the Slingshot^35^ package (v2.10.0). SCTransform-corrected counts and previously generated PCA embeddings were used for trajectory inference. Since no naïve population was present in the transition of interest, the root was not specified. Gene expression changes along pseudotime were analyzed using generalized additive models, specifically by fitting the formula z ∼ lo(t) in **R**, where *z* represents the expression of each gene and *t* denotes the Slingshot pseudotime. This model uses a loess smoother to capture gene expression trends over time. Genes with significant expression changes along pseudotime (*p* < 0.05) were selected, with the top 50 non-ribosomal and non-mitochondrial genes (ranked by *p*-values, ascending order) highlighted. Standardized expression levels were visualized in heatmaps to illustrate gene expression patterns over pseudotime.

### Geneformer

To identify genes whose overexpression in T cells from minimally expanded clones (top 100+ clones by size) could shift their state toward those from highly expanded clones (top 1-10 clones), we leveraged Geneformer^48^ (v0.1.0). For donor-site-specific subtypes (excluding peripheral blood, PB) that showed significant differential expression between these groups, we tokenized cells using Geneformer’s TranscriptomeTokenizer, retaining protein-coding and miRNA genes by default. The initial state (cells from the top 100+ clones) and the goal state (cells from the top 1-10 clones) were represented by the exact mean [CLS] token embeddings of cells in each group. Geneformer simulated gene overexpression by reordering the target gene’s token to the first position within the input sequence, which contained all gene tokens of a cell. Since gene token positions are determined by expression rank, this is functionally equivalent to promoting the overexpressed gene’s rank to 1 in the input. Perturbed embeddings were then generated via Geneformer, and the directional cosine similarity shift toward the goal state was quantified. Specifically, Geneformer defines the cell state cosine shift as cos(perturbed cell embeddings, goal state embeddings) - cos(cell embeddings, goal state embeddings). To ensure robustness, we perturbed up to 1,000 cells per gene. Significance was assessed using Wilcoxon rank-sum tests, comparing shifts induced by each gene against the background distribution of all perturbations, with Benjamini-Hochberg correction. Genes with adjusted *p* < 0.05 and a mean cosine shift > 0.002 (empirically selected to reduce background noise) were prioritized.

## Data Availability

Annotated data for this study can be obtained at https://zenodo.org/records/15002914.

## Code Availability

Scripts used for this analysis can be accessed at https://github.com/Brubaker-Lab/gut-liver-TRM.

## Acknowledgements

Single-cell sequencing was performed at the JHU Single Cell and Transcriptomics Core. The study was supported by grants from NIGMS 5R35GM146900. R. R. and D. K. B are supported by an award from the Good Ventures Foundation and Open Philanthropy, as well as start-up funds from Case Western Reserve University and University Hospitals.

## Author Contributions

R. R. analyzed data, made figures, and wrote the manuscript. M. U. procured donor tissues, performed experiments, prepared samples for single-cell TCR and RNA sequencing, and wrote the manuscript. M. F. S. performed experiments. D. K. B. wrote and edited the manuscript. M. T. designed experiments, coordinated the project, and wrote and edited the manuscript.

