## Supplementary figures and images for "Single-Cell Analysis Reveals Tissue-Specific T Cell Adaptation and Clonal Distribution Across the Human Gut-Liver-Blood Axis"

### Supplemental Figure 1

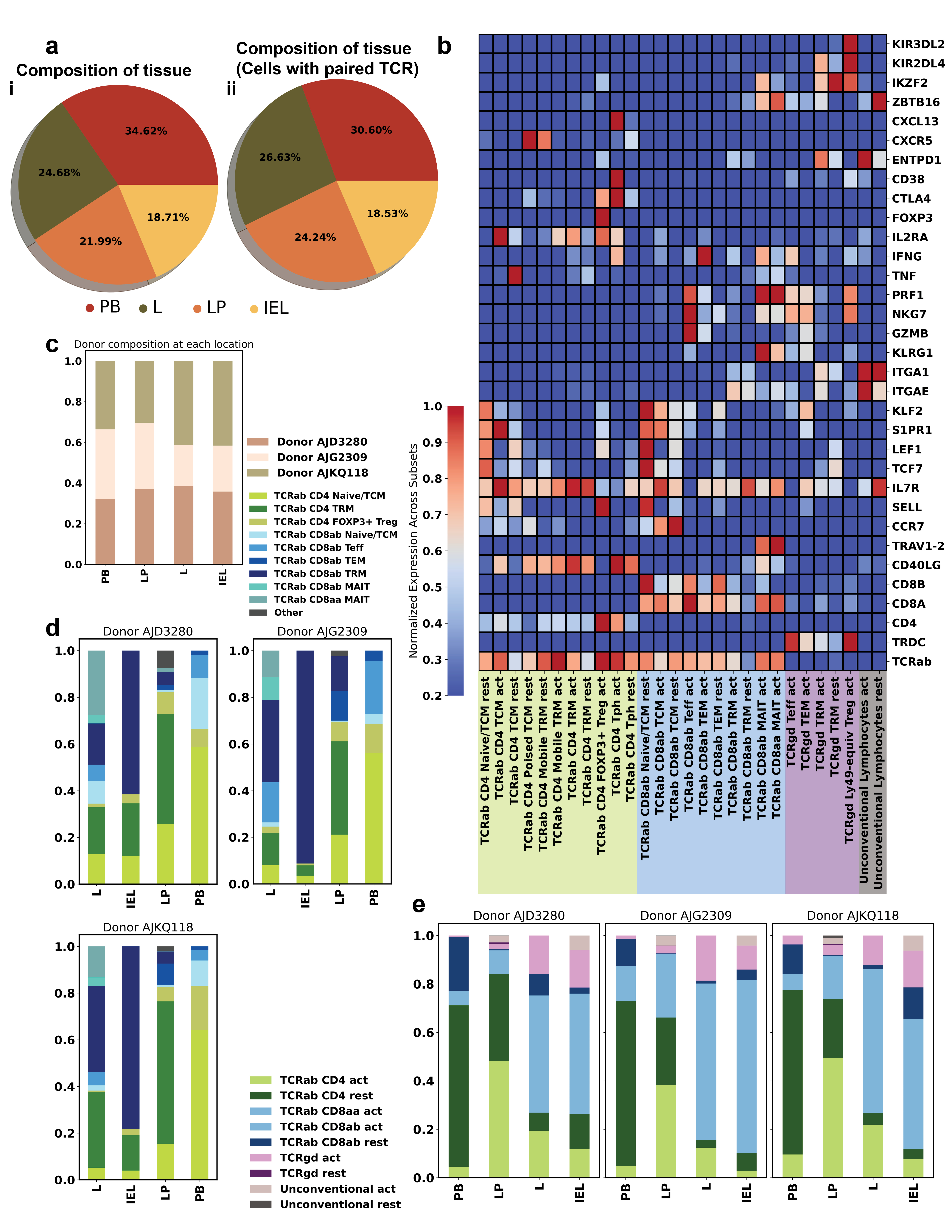

### Supplemental Figure 2

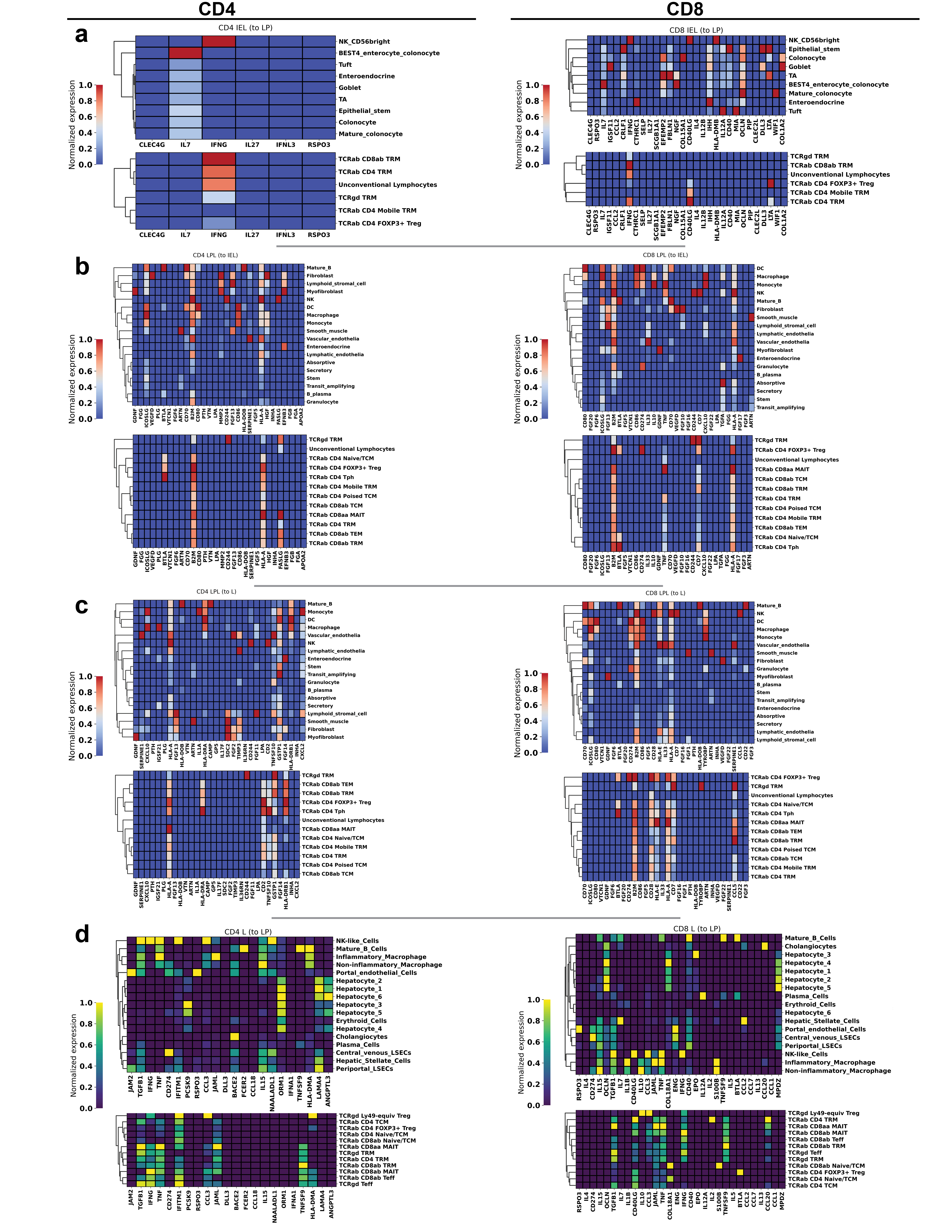

### Supplemental Figure 3

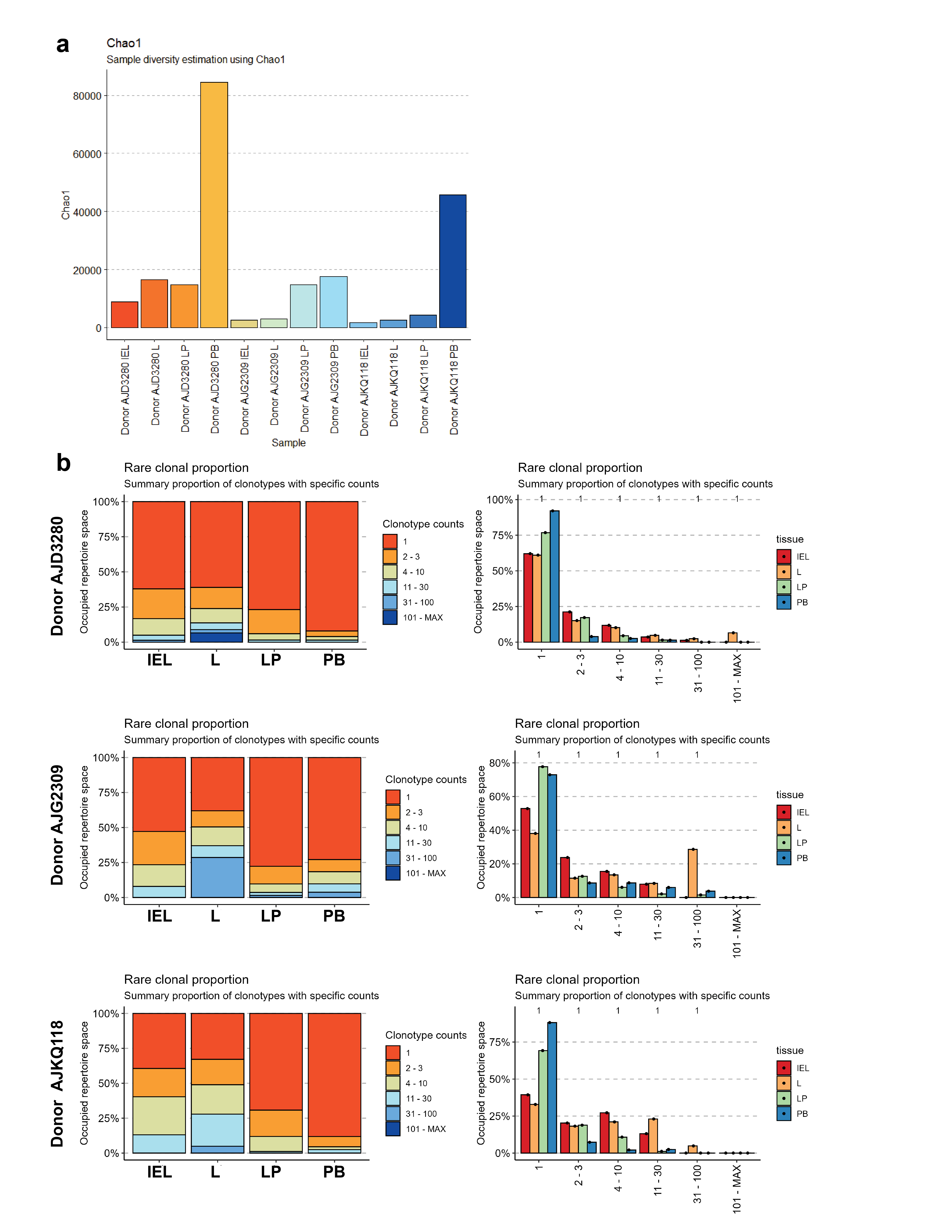

### Supplemental Figure 4

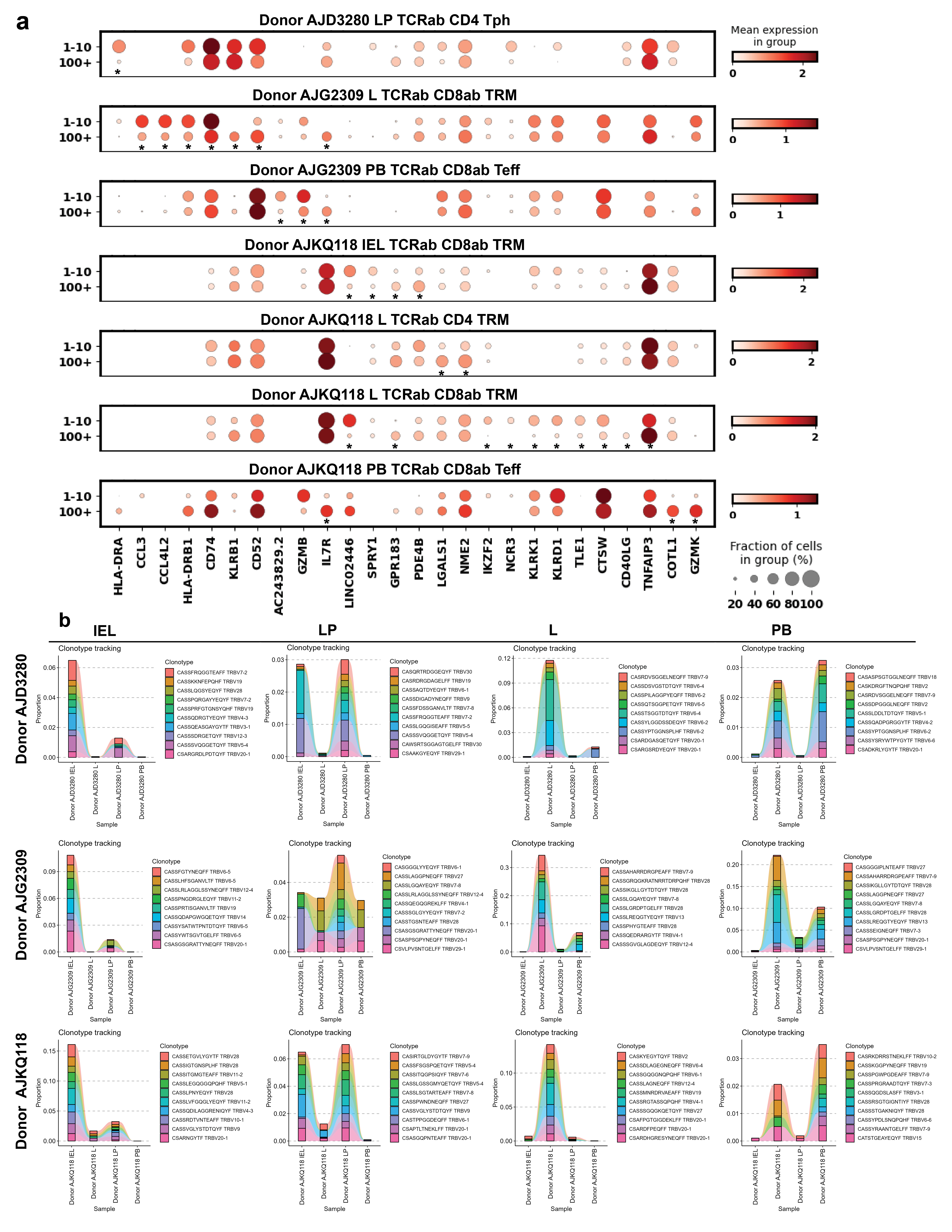

### Supplemental Figure 5

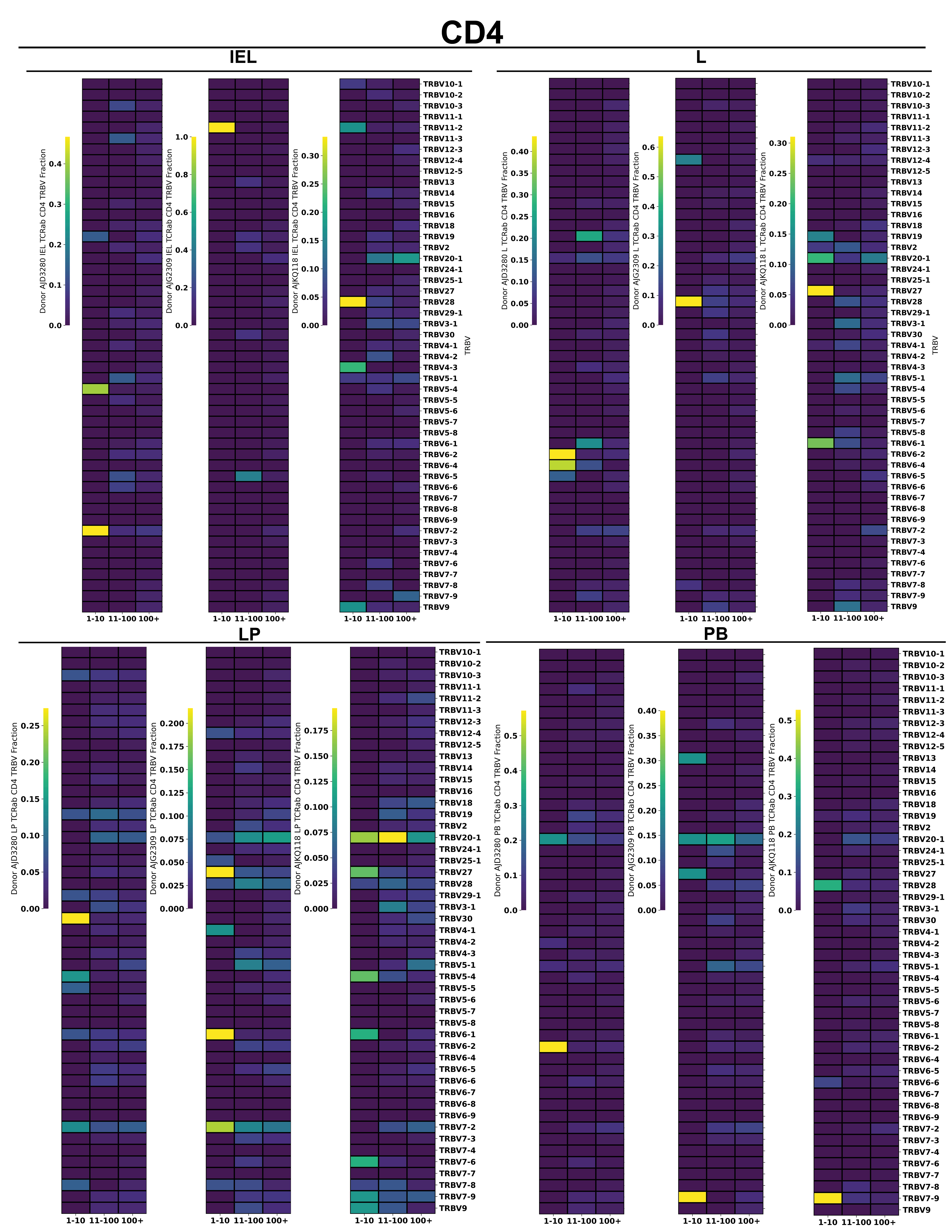

### Supplemental Figure 6

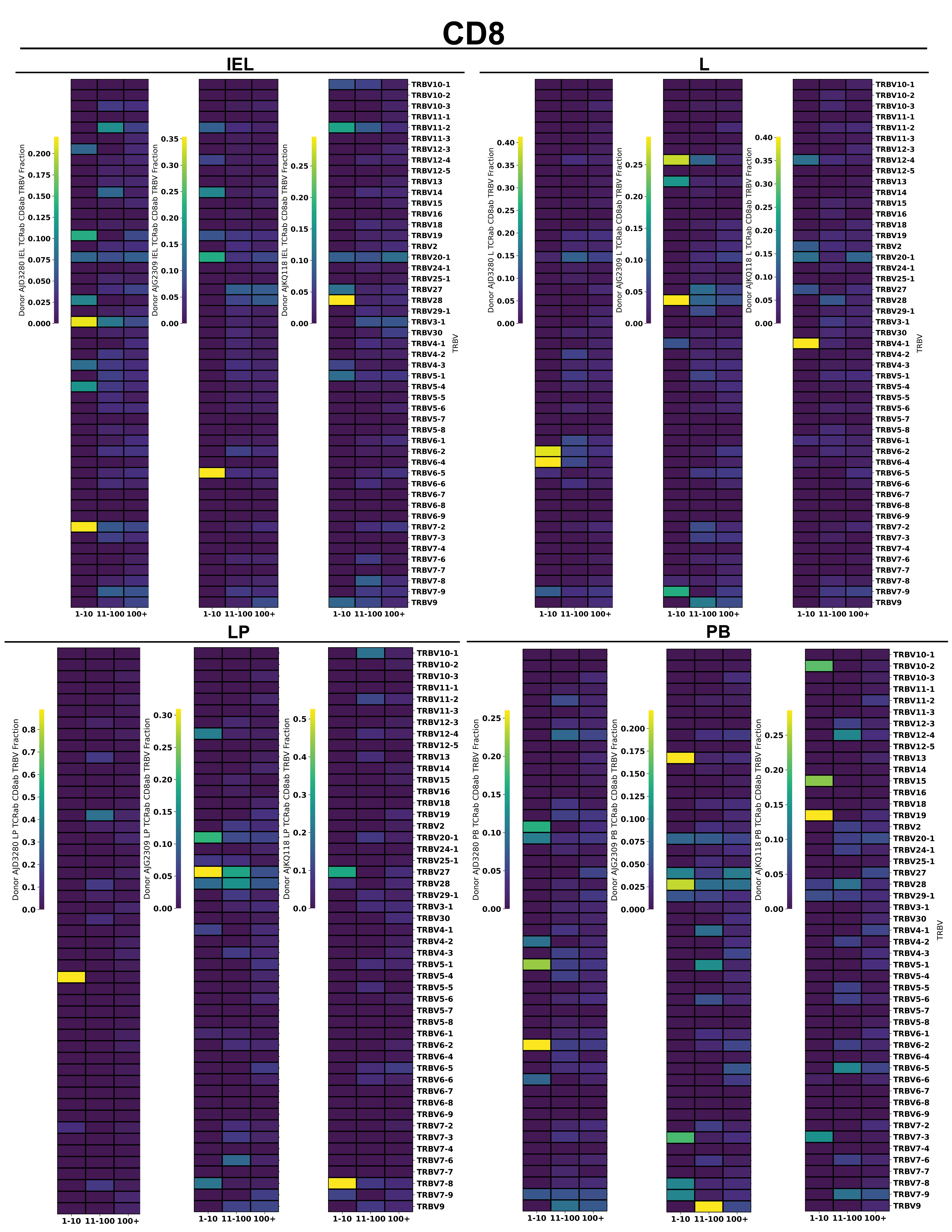
